# openTSNE: a modular Python library for t-SNE dimensionality reduction and embedding

**DOI:** 10.1101/731877

**Authors:** Pavlin G. Poličar, Martin Stražar, Blaž Zupan

## Abstract

**Summary:** Point-based visualisations of large, multi-dimensional data from molecular biology can reveal meaningful clusters. One of the most popular techniques to construct such visualisations is t-distributed stochastic neighbor embedding (t-SNE), for which a number of extensions have recently been proposed to address issues of scalability and the quality of the resulting visualisations. We introduce openTSNE, a modular Python library that implements the core t-SNE algorithm and its extensions. The library is orders of magnitude faster than existing popular implementations, including those from scikit-learn. Unique to openTSNE is also the mapping of new data to existing embeddings, which can surprisingly assist in solving batch effects.

**Availability:** openTSNE is available at https://github.com/pavlin-policar/openTSNE.

---

The abundance of high-dimensional data sets in molecular biology calls for techniques for dimensionality reduction, and in particular for methods that can help in the construction of data visualizations. Popular approaches for dimensionality reduction include principal component analysis, multidimensional scaling, and uniform manifold approximation and projections (1). Among these, t-distributed stochastic neighbor embedding (t-SNE) (2) lately received much attention as it can address high volumes of data and reveal meaningful clustering structure. Most of the recent reports on singlecell gene expression data start with an overview of the cell landscape, where t-SNE embeds high-dimensional expression profiles into a two-dimensional space (3, 4). Fig. 1.a presents an example of one such embedding.

**Fig. 1.**
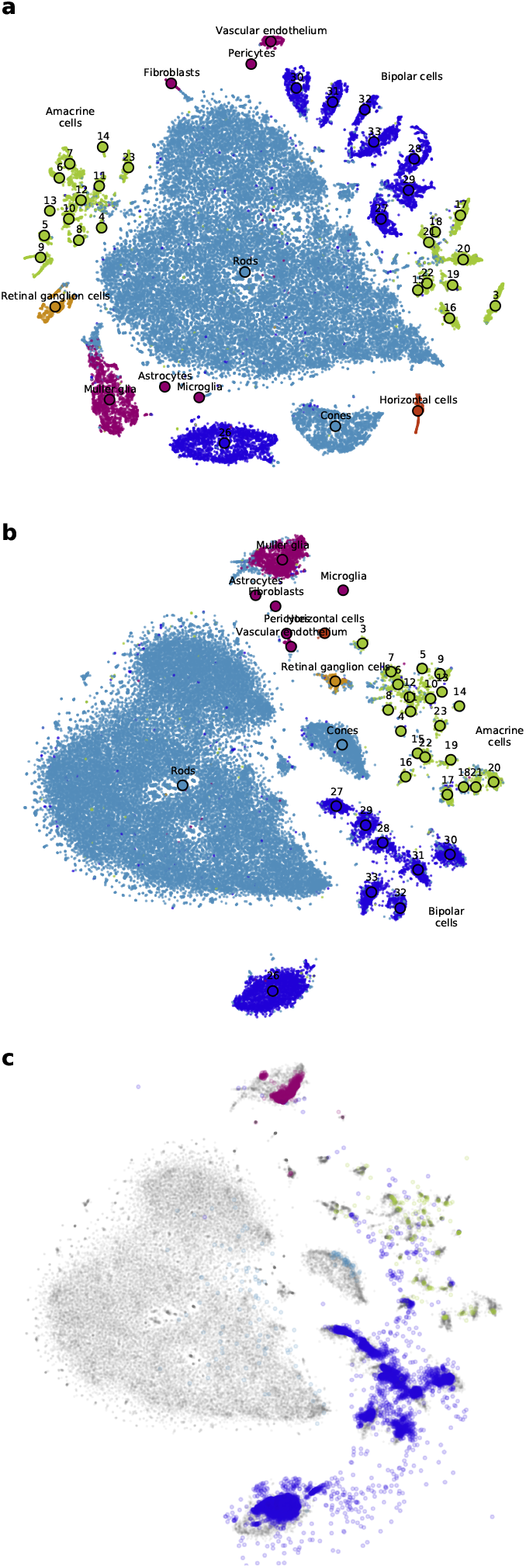
Example application of `openTSNE` on mouse retina cells from Macosko et al. (3) and Shekhar et al. (4). a) Standard t-SNE embedding using random initialization and perplexity of 30. b) t-SNE embedding using multi-scale affinities leads to better global cluster organization. Cluster annotations in both a) and b) are from Macosko *et al.* c) Embedding of independent mouse retinal cells from Shekhar *et al.* (points in color) to a reference t-SNE visualization of data from Macosko *et al.* (points in gray) places cells in the clusters that match classifications from original publications and alleviates batch effects.

Despite its utility, t-SNE has been criticized for poor scalability when addressing large data sets, lack of global organization *t-SNE focuses on local clusters that are arbitrarily scattered in the low-dimensional space* and absence of theoretically-founded implementations to map new data into existing embeddings (5, 6). Most of these shortcomings, have recently been addressed. Linderman et al. (7) sped-up the method through an interpolation-based approximation, reducing the time complexity to be merely linear dependent on the number of samples. Kobak and Berens (8) proposed several techniques to improve global positioning, including estimating similarities with a mixture of Gaussian kernels. While no current popular software library supports mapping of new data into reference embedding, van der Maaten (9) proposed a related approach using neural networks.

We introduce `openTSNE`, a comprehensive Python library that implements t-SNE and all its recently proposed extensions. The library is compatible with the Python data science ecosystem (e.g., `numpy, sklearn, scanpy`). Its modular design encourages extendibility and experimentation with various settings and changes in the analysis pipeline. For example, the following code uses multiscale similarity kernels to construct the embedding from Fig. 1.b.

~~~
adata =
   anndata.read_h5ad(“macosko_2015.h5ad”)
affinities = openTSNE.affinity.Multiscale(
  adata.obsm[“pca”], perplexities=[50,
     500], metric=“cosine”)
init = openTSNE.initialization.pca(
 adata.obsm[“pca”])
embedding = TSNEEmbedding(init, affinities)
embedding.optimize(n_iter=250,
    exaggeration=12, momentum=0.5,
    inplace=True)
embedding.optimize(n_iter=750,
    momentum=0.8, inplace=True)
~~~

Here, we first read the data, define the affinity model based on two Gaussian kernels with varying perplexity, use a PCA-based initialization, and run the typical two-stage t-SNE optimization. Notice that the code for the standard t-SNE used for Fig. 1.a is similar but uses only a single kernel (`perplexities=[30]`).

The proposed `openTSNE` library is currently the only Python t-SNE implementation that supports adding new samples into constructed embedding. For example, we can reuse the `embedding` created above to map new data into existing embedding space in the following code,

~~~
data =
   anndata.read_h5ad(“shekhar_2016.h5ad”)
adata, data = find_shared_genes(adata, data)
genes = select_genes(adata.X, n=1000)
embedding.affinities =
   affinity.PerplexityBasedNN(
  adata[:, genes].X, perplexity=30,
     metric=“cosine”)
new_embedding = embedding.transform(data[:,
    genes].X)
~~~

which loads and prepare the new data, defines the affinity model, and computes the embeddings. `openTSNE` embeds new data points independently of one another, without changing the reference embedding. The example of combining two data sets with mouse retina cells is shown on Fig. 1.c, where the cells from the secondary data set are matched to the cells in the reference embedding. This procedure can therefore be used to handle batch effects (10), a key obstacle in molecular biology when dealing with the data from different sources (11). The code used to plot the embeddings is not shown for brevity, but is, together with other examples, available on `openTSNE`’s GitHub page.

Our Python implementation introduces computational overhead: `openTSNE` is about 25% slower than `FIt-SNE` (7), a recent t-SNE implementation in C++. However, openTSNE is still orders of magnitude faster than other Python implementations, including those from `scikit-learn` and `MulticoreTSNE` (see Benchmarks in `openTSNE` documentation on GitHub). An example data set with 200,000 cells is processed in more than 90 minutes with scikit-learn and in less than 4 minutes with `openTSNE`. The framework includes controlled execution (callback-based progress monitoring and control), making it suitable for interactive data exploration environments such as Orange (12). A pure Python implementation offers distinct advantages that include integration with Python’s rich data science infrastructure and ease of installation through PyPI and *conda*.

## Funding and Acknowledgements

The support for this work was provided by the Slovenian Research Agency Program Grant P2-0209, and by European Regional Development Fund and the Slovenian Ministry of Education through BioPharm project. We want to thank George Linderman and Dmitry Kobak for helpful discussions on extensions of t-SNE.

